# Regulation Mechanisms of the Dual ATPase in KaiC

**DOI:** 10.1101/2021.10.28.466029

**Authors:** Yoshihiko Furuike, Atsushi Mukaiyama, Shinichi Koda, Damien Simon, Dongyan Ouyang, Kumiko Ito-Miwa, Shinji Saito, Eiki Yamashita, Taeko Nishiwaki-Ohkawa, Kazuki Terauchi, Takao Kondo, Shuji Akiyama

## Abstract

KaiC is a dual ATPase, with one active site in its N-terminal domain and another in its C-terminal domain, that drives the circadian clock system of cyanobacteria through sophisticated coordination of the two sites. To elucidate the coordination mechanism, we studied the contribution of the dual ATPase activities in the ring-shaped KaiC hexamer and these structural bases for activation and inactivation. At the N-terminal active site, a lytic water molecule is sequestered between the N-terminal domains, and its reactivity to ATP is controlled by the quaternary structure of the N-terminal ring. The C-terminal ATPase activity is regulated mostly by water-incorporating voids between the C-terminal domains, and the size of these voids is sensitive to phosphoryl modification of S431. The N-terminal ATPase activity inversely correlates with the affinity of KaiC for KaiB, a clock protein constitutes the circadian oscillator together with KaiC and KaiA, and the complete dissociation of KaiB from KaiC requires KaiA-assisted activation of the dual ATPase. Delicate interactions between the N-terminal and C-terminal rings make it possible for the components of the dual ATPase to work together, thereby driving the assembly and disassembly cycle of KaiA and KaiB.

**Significance Statement:** KaiC, a core clock protein in the cyanobacterial circadian clock system, hydrolyzes ATP at two distinct sites in a slow but ordered manner to measure the circadian time scale. We used biochemical and structural biology techniques to characterize the properties and interplay of dual ATPase active sites. Our results show that the N-terminal and C-terminal ATPases communicate with each other through an interface between the N-terminal and C-terminal domains in KaiC. The dual ATPase sites are regulated rhythmically in a concerted or opposing manner dependent on the phase of the circadian clock system, controlling the affinities of KaiC for other clock proteins, KaiA and KaiB.

## Introduction

The circadian clock is an endogenous timekeeping system that rhythmically controls a variety of biological phenomena with a period of about 24 hours. The rhythms of the circadian clock remain stable without external stimuli even under homeostatic conditions (self-sustained oscillation), and their period is kept constant even when the ambient temperature changes (temperature compensation). Moreover, the circadian clock system responds by shifting its internal phase back and forth depending on external stimuli such as light and temperature, and through this phase response, the internal phase of the timing system can be synchronized with the external phase of the environmental cycle (synchronization). These unique properties allow organisms to adapt to the daily environmental cycles on Earth.

KaiC is a core clock protein that regulates the circadian rhythm of the cyanobacterium *Synechococcus elongatus* PCC 7942 (1). This protein utilizes two types of ATP molecules to produce diverse chemical reactions and thereby generate a temperature-compensated circadian rhythm (2-4). ATP molecules bind to a Walker motif present in two tandemly replicated domains, specifically N-terminal C1 and C-terminal C2 domains (Fig. 1*A*). The binding of ATP triggers the oligomerization of KaiC into a double-ring hexamer (5) (Fig. 1*A*). The ATP molecule bound to the C2 domain (C2-ATP) is the primary source of a phosphoryl group that is transferred to S431 and T432 of KaiC (autokinase) and then removed (autophosphatase) (6, 7). In the presence of KaiA and KaiB, the state of the phosphorylation sites changes periodically even in vitro (6-8): ST → SpT → pSpT → pST → ST, where S, T, pS, and pT represent S431, T432, phosphorylated S431, and phosphorylated T432, respectively. In addition, it has been suggested that a small amount of C2-ATP is hydrolyzed into C2-ADP (C2-ATPase) via a complicated reaction scheme (9, 10). On the other hand, the frequency of the phosphorylation cycle is proportional to the hydrolysis (C1-ATPase) rate of C1-bound ATP (C1-ATP) in the absence of KaiA and KaiB (11, 12). The C1-ATPase activity greatly exceeds the C2-ATPase activity under steady state and is temperature compensated (12). Overall ATPase activities, including those of C1 and C2 (C1/C2-ATPase), are known to oscillate with the circadian period (12) in the presence of KaiA and KaiB (Fig. 1*B*), and also to reveal a damping oscillatory relaxation even without KaiA and KaiB (11).

**Figure 1.**
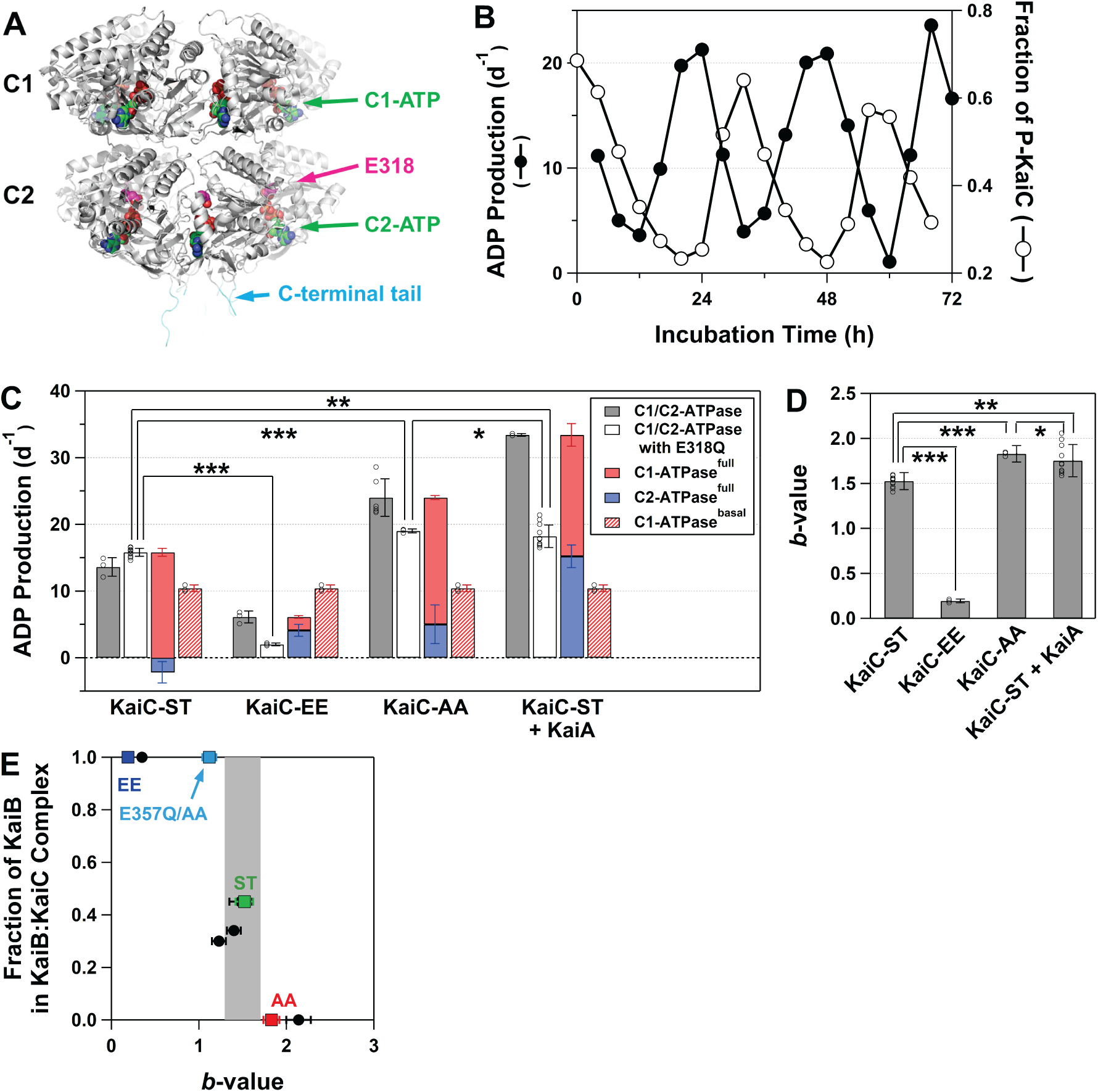
Regulation of the dual C1/C2-ATPase in KaiC. (*A*) Active sites of the dual C1/C2-ATPase in the KaiC hexamer (5). (*B*) Rhythmic changes in C1/C2-ATPase activity of KaiC (*left*) and the fraction of phosphorylated KaiC (P-KaiC, *right*) in the presence of KaiA and KaiB at 30°C. (*C*) Steady-state rates of ADP production. Gray, white, and hatched bars represent the ATPase activities of full-length KaiCs (C1/C2-ATPase; contributions from C1 and C2), full-length KaiCs carrying the E318Q substitution (C1-ATPase^full^; contribution from C1), and truncated KaiCs consisting solely of the C1 domain (C1-ATPase^basal^; C1 basal activity without the C2 domains). Red and blue bars indicate C1-ATPase^full^ and C2-ATPase^full^ (C1/C2-ATPase minus C1-ATPase^full^) activities, respectively, in the full-length KaiCs. ATPase activities were assayed by measuring ADP production for 10–16 h. In the case of KaiC-ST (0.20 mg/mL) + KaiA (0.12 mg/mL), data are given as apparent activities estimated by the ADP production for 10 h. The ATPase activities were analyzed using one-way ANOVA (F_3,26_ = 2.975, p = 3.95×10^−17^) and Bonferroni post hoc *t* test: ***significant (p < 0.0005 / 6), **significant (p < 0.005 / 6), *insignificant (p > 0.05 / 6). (*D*) *b*-values defined as the ratio of C1-ATPase^full^ (red bars in panel C) to C1-ATPase^basal^ (hatched bars in panel C) (details in Materials and Methods). The *b*-values were analyzed using one-way ANOVA (F_3,26_ = 2.975, p = 3.95×10^−17^) and Bonferroni post hoc *t* test: ***significant (p < 0.0005 / 6), **significant (p < 0.005 / 6), *insignificant (p > 0.05 / 6). (*E*) Potential correlation between *b*-value and KaiB–KaiC affinity. The fraction of KaiB forming KaiB–KaiC complexes was determined by native PAGE analysis (*SI Appendix*, Fig. S1*A*). C1/C2-ATPase and *b*-values for a series of mutants (black circles) are compiled in *SI Appendix*, Table S3.

The C1/C2-ATPase of KaiC is an important research target to achieve a better understanding of the circadian clock system in cyanobacteria (3), as it closely relates to oscillatory, period-tuning, and temperature-compensating properties. Although much effort has been devoted to the biochemical (12-18) and structural analyses (11, 19-21) of KaiC ATPase, the mechanisms of its activation and inactivation remain unknown. In this study, we performed a biochemical analysis of C1/C2-ATPase and a crystal structure analysis of the activated and inactivated forms of KaiC. Our results suggest that the components of the dual ATPase communicate with each other through C1–C2 interaction to control the assembly and disassembly cycle of KaiA and KaiB.

## Results

### Contributions of C1- and C2-ATPases

The steady-state C1/C2-ATPase activity of fully dephosphorylated KaiC (KaiC-ST) was 13.6 ± 1.4 d^-1^ at 30°C (gray bars in Fig. 1*C*). To estimate the contribution of C1-ATPase in full-length KaiC (C1-ATPase^full^), we inactivated the kinase and C2-ATPase by replacing the catalytic glutamate 318 with glutamine (13-16, 22). The resultant KaiC^E318Q^-ST exhibited C1-ATPase^full^ activity of 15.8 ± 0.6 d^-1^ (white bars in Fig. 1*C*), which was comparable to the C1/C2-ATPase activity of KaiC-ST. Given that this estimated activity of KaiC^E318Q^-ST reflects the C1-ATPase^full^ activity of KaiC-ST (red bars in Fig. 1*C*), the contribution of C2-ATPase^full^ can be estimated as the difference between C1/C2-ATPase and C1-ATPase^full^, which was as small as −2.2 ± 1.6 d^-1^ (blue bars in Fig. 1*C*). On the other hand, the ATPase activity of a truncated KaiC (KaiC1) consisting solely of the C1 domain (11), which reflects the basal activity of the C1-ATPase (C1-ATPase^basal^) without the influence of the C2 domain, was 10.4 ± 0.5 d^-1^ (hatched bars in Fig. 1*C*). The *b*-value, defined as the ratio of C1-ATPase^full^ to C1-ATPase^basal^, was 1.52 ± 0.10 for KaiC-ST (Fig. 1*D*), reflecting the upregulation of its C1-ATPase by the C2 domain. These results suggest that the C1-ATPase^full^ activity in KaiC-ST is ∼50% upregulated by the C2 domain (Fig. 1*D*), while that of the C2-ATPase^full^ is negligibly small (Fig. 1*C*).

To investigate how phosphoryl modifications affected C1/C2-ATPase, we measured the C1/C2-ATPase activity of a KaiC-pSpT–mimicking S431E/T432E KaiC mutant (KaiC-EE). The C1-ATPase^full^ and C2-ATPase^full^ activities of KaiC-EE were 2.0 ± 0.2 and 4.1 ± 0.9 d^-1^, respectively (Fig. 1*C*). The *b*-value of KaiC-EE was estimated to be 0.19 ± 0.02, indicating 80% downregulation of its C1-ATPase^full^ (Fig. 1*D*). By contrast, an S431A/T432A KaiC mutant (KaiC-AA), which may be treated as a phospho-mimic mutant of KaiC-ST, exhibited activation of the C1/C2-ATPase (24.0 ± 2.8 d^-1^) as reported previously (12). While the C2-ATPase^full^ activity of KaiC-AA was 5.0 ± 2.9 d^-1^, which was comparable to that of KaiC-EE (4.1 ± 0.9 d^-1^) (Fig. 1*C*), the C1-ATPase^full^ activity was 80% upregulated as high as 19.0 ± 0.3 d^-1^ (*b* = 1.83 ± 0.09 in Fig. 1*D*). The excessive activation of C1/C2-ATPase in KaiC-AA relative to KaiC-ST indicates that KaiC-AA is not an exact functional counterpart of KaiC-ST.

We found that KaiC-AA is rather similar to a KaiA-stimulated state of KaiC-ST. KaiA was previously shown to activate the ATPase activity of KaiC (12). Consistent with that study, the addition of KaiA to KaiC-ST resulted in the activation of the dual ATPase. The C1-ATPase^full^ activity of KaiA-stimulated KaiC-ST was 18.2 ± 1.7 d^-1^ (Fig. 1*C*), and was ∼80% upregulated relative to KaiC-ST alone (*b* = 1.75 ± 0.18 in Fig. 1*D*) (*t* test: **significant; p < 0.005 / 6, Fig. 1*C*). The contribution of C2-ATPase^full^ in KaiC-ST became very obvious in the presence of KaiA, with enhanced activity of 15.2 ± 1.7 d^-1^ for the initial 10 h (Fig. 1*C*). These observations suggest that the KaiA-stimulation of C1/C2-ATPase of KaiC-ST is mimicked in KaiC-AA even without KaiA.

### Structural Basis for Up- and Downregulation of C1-ATPase^full^

ATP hydrolysis includes a reaction step in which a lytic water molecule (W1) attacks the phosphorus atom (P_γ_) of the terminal gamma-phosphate of ATP. Thus, the relative positioning of W1 and P_γ_ is one of the crucial factors determining the reaction efficiency, as reported previously (11). To investigate the W1 position, we sought to crystallize KaiC-AA (*SI Appendix*, Table S1) and to compare the results with those for KaiC-ST and KaiC-pSpT, which we previously reported (23). We could identify a reasonable electron density for W1 in all cases (Fig. 2, *A*–*C* and *SI Appendix*, Table S2) except for KaiC-pSpT.

**Figure 2.**
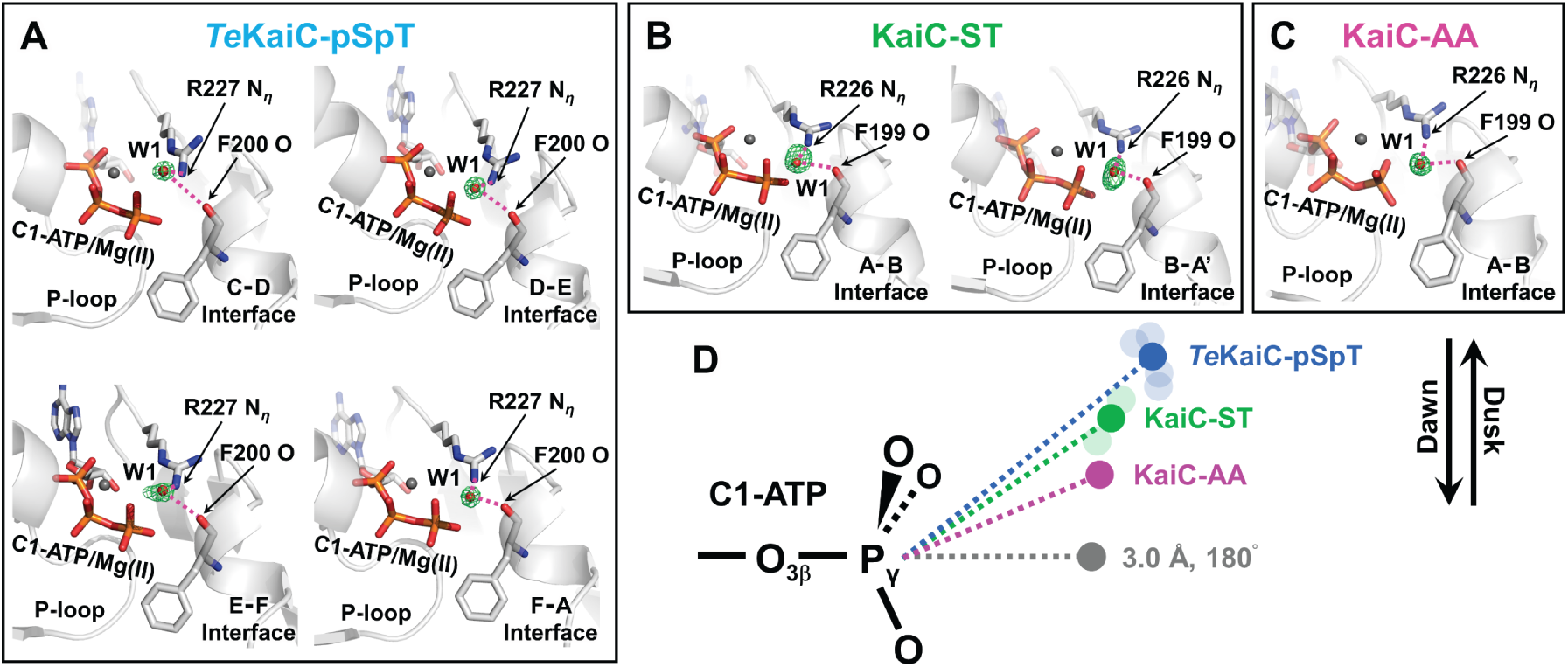
Active-site structures of C1-ATPase in KaiC. Lytic water molecules (W1: red spheres) identified for (*A*) *Te*KaiC-pSpT, (*B*) KaiC-ST, and (*C*) KaiC-AA. *F*o–*F*c omit maps (green mesh) in panels *A*–*C* are contoured respectively as follows: 3.0σ, 5.0σ, 2.8σ, and 2.6σ; 3.3σ and 2.0σ; and 4.0σ. W1 are sequestered through hydrogen bonds (magenta dotted lines) by F199 and R226 (F200 and R227 in *Te*KaiC) to regulate their position relative to the ATP molecule bound to the C1 domain. (*D*) Schematic drawing of dawn- and dusk-phase repositioning of W1 (*SI Appendix*, Table S2). Dark-colored circles correspond to the mean W1 positions from independent C1–C1 interfaces (pale-colored circles).

Fortunately, however, another 2.2-Å crystal structure of thermophilic KaiC from *Thermosynechococcus elongatus* (*Te*KaiC) was determined in the *Te*KaiC-pSpT state (*SI Appendix*, Table S1), with a clear electron density for W1.

In *Te*KaiC-pSpT (Fig. 2*A*), W1 was hydrogen-bonded to the carbonyl oxygen atom of F200 and R227 N_η_ (F199 and R226 in KaiC) and sequestered at the position most unfavorable for S_N_2-type ATP hydrolysis, with a W1–P_γ_ distance of 5.1 Å and a W1–P_γ_–O_3β_ angle of 141° (Fig. 2*D*). By contrast, W1 was located at a less unfavorable position in KaiC-ST (Fig. 2*B*), with a shorter W1–P_γ_ distance (4.0 Å) and a larger W1–P_γ_–O_3β_ angle (147°) (Fig. 2*D*). W1 was found in positions more suitable for ATP hydrolysis in KaiC-AA (Fig. 2, *C* and *D*). The descending and ascending orders of the W1–P_γ_ distance and the W1–P_γ_–O_3β_ angle (Fig. 2*D*), respectively, matched the increasing order of the C1-ATPase^full^ activity (KaiC-EE(pSpT) < KaiC-ST < KaiC-AA, Fig. 1, *C* and *D*). These results confirm that the relative positioning of W1 and P_γ_ is one of the factors impacting up- and downregulation of C1-ATPase^full^.

### Structural Basis for Up- and Downregulation of C2-ATPase^full^

The electron densities for lytic water molecules involved in C2-ATPase could not be identified in the crystal structures of KaiC-pSpT, *Te*KaiC-pSpT, KaiC-ST, or KaiC-AA, possibly due to their dynamic nature. On the other hand, we found unique variation patterns in a tunnel-shaped void in the C2–C2 interface (Fig. 3), through which bulk water molecules migrate to the C2-ATPase active site.

**Figure 3.**
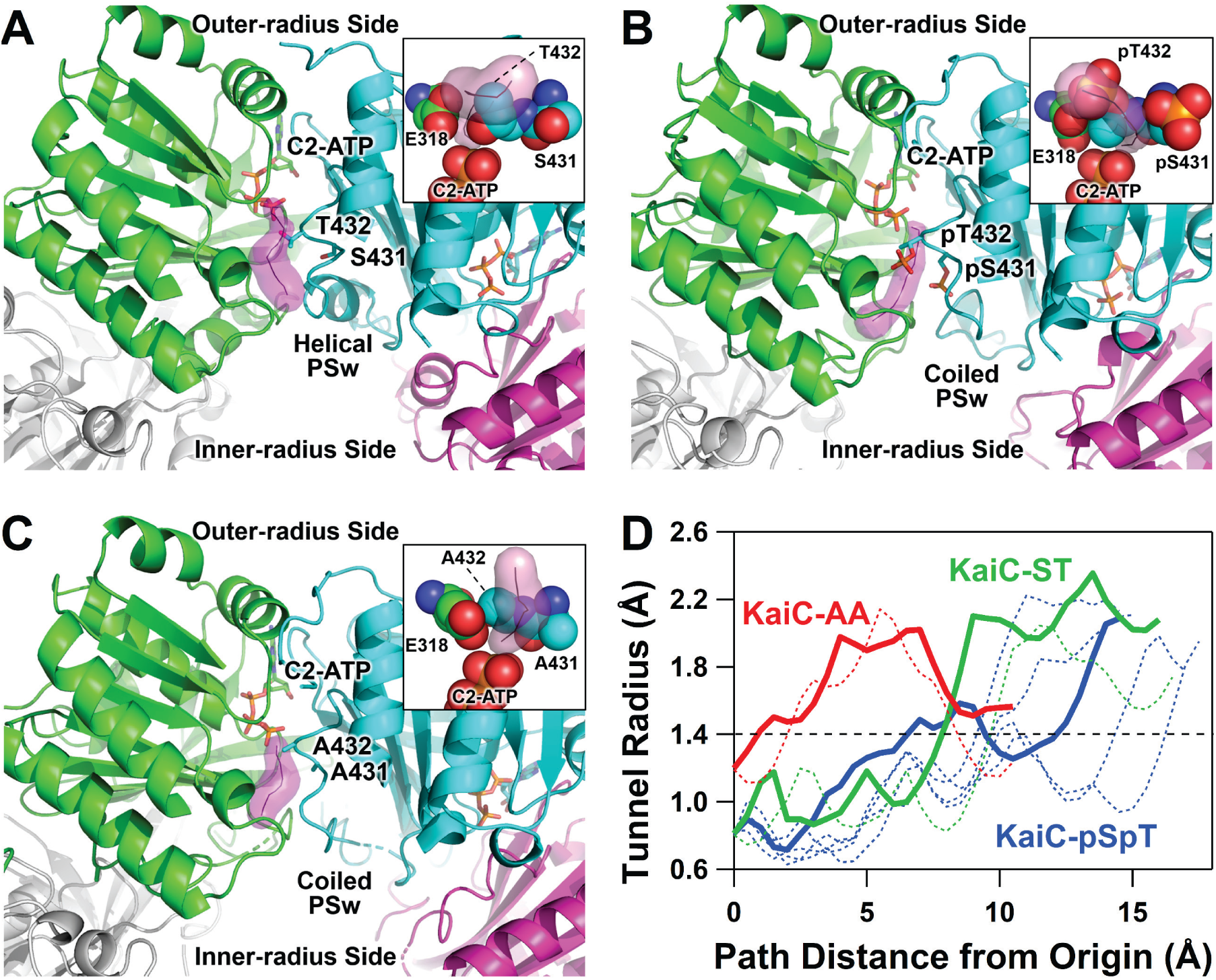
Tunnel-shaped voids extending along C2–C2 interfaces from C2-ATP to the inner-radius side of the hexamer. Zoomed-in view of C2–C2 interfaces for (*A*) KaiC-ST, (*B*) KaiC-pSpT, and (*C*) KaiC-AA. Tunnels (magenta surface) extend from the vicinity of P_γ_ of C2-ATP to the inner-radius side of the hexamer. The insets depict representative amino acids, shown as space-filling models, located on the distal side of the γ-phosphate of C2-ATP. (*D*) Radius of the tunnels as a function of path distance from the origin (near P_γ_ of C2-ATP). The shortest and widest tunnels in KaiC-ST (green), KaiC-pSpT (blue), and KaiC-AA (red) are highlighted by thick lines. In KaiC-ST, the first half of the tunnel is undulated (thick black line in panel A) and then narrowed by bottleneck residues down to a radius of ∼0.8 Å, far smaller than the radius of a water molecule (1.4 Å, dotted line in the panel D). Thus, the tunnel of KaiC-ST is not wide enough to freely take up water molecules from the inner-radius side. By contrast, the tunnel without a bottleneck that exhibits straight extension in KaiC-AA is ready to transfer water molecules from the inner-radius side to the active site of C2-ATPase.

In KaiC-ST, the distal side of the γ-phosphate of C2-ATP is stereochemically crowded by several residues (E318, S431, and T432) and consequently does not have enough space to stably accommodate any water molecules (inset of Fig. 3*A*). Although a tunnel-shaped void was confirmed near the distal side, it was undulated (Fig. 3*A*) and too narrow (Fig. 3*D*) to freely take up water molecules (radius of ∼1.4 Å) from the inner-radius side of the hexamer. The corresponding water tunnels were not uniform within the KaiC-pSpT hexamer, but the shortest one suggests more effective uptake by KaiC-pSpT than by KaiC-ST (Fig. 3, *B* and *D*). By contrast, the distal side of KaiC-AA was more relaxed. A wider straight channel (Fig. 3*C*) connected the space in the distal side to the exterior of the hexamer, and appeared to allow effective migration of water molecules into the active site of C2-ATPase (Fig. 3*D*). Furthermore, the side chain of catalytic E318 was positioned at one end of the water tunnel and could immediately activate the migrated water molecules (inset of Fig. 3*C*). The orders of the shortness and width of the tunnel-shaped voids approximately matched the increasing order of the C2-ATPase^full^ activity (KaiC-ST < KaiC-EE(pSpT) < KaiC-AA, Fig. 1*C*). These observations provide a structural explanation for why C2-ATPase is downregulated in KaiC-ST but upregulated in KaiC-EE and KaiC-AA.

### Quaternary Structural Changes

A region upstream of the dual phosphorylation sites in the C2 domain called phosphor-switch (PSw) (23) adopts coiled and helical conformations in KaiC-pSpT (EE) and KaiC-ST, respectively (Fig. 4*A*). Our previous study (23) suggested that this fold switch of PSw influences the quaternary structure of the C1 ring through the C1–C2 interface. When superimposed using the C1 domains of KaiC-pSpT and KaiC-ST as references (Fig. 4*C*), one of the two adjacent C1 domains (*right subunit* in Fig. 4C) of KaiC-pSpT shifted by ∼1 Å away from the C2 domain, but conversely the other C1 domain (*left subunit* in Fig. 4C) shifted towards the C2 domain. These observations indicate that during the helix-to-coil transition of PSw from KaiC-ST to KaiC-pSpT, every C1 domain gently tilts and then leans over a neighboring C1 domain (Fig. 4*C*). This tilting and sliding movement causes W1, which is sequestered by F199 and R226 (F200 and R227 in *Te*KaiC), to move one position further toward the ATP molecule bound to the C1 domain (Fig. 2*D* and Fig. 4*E*).

**Figure 4.**
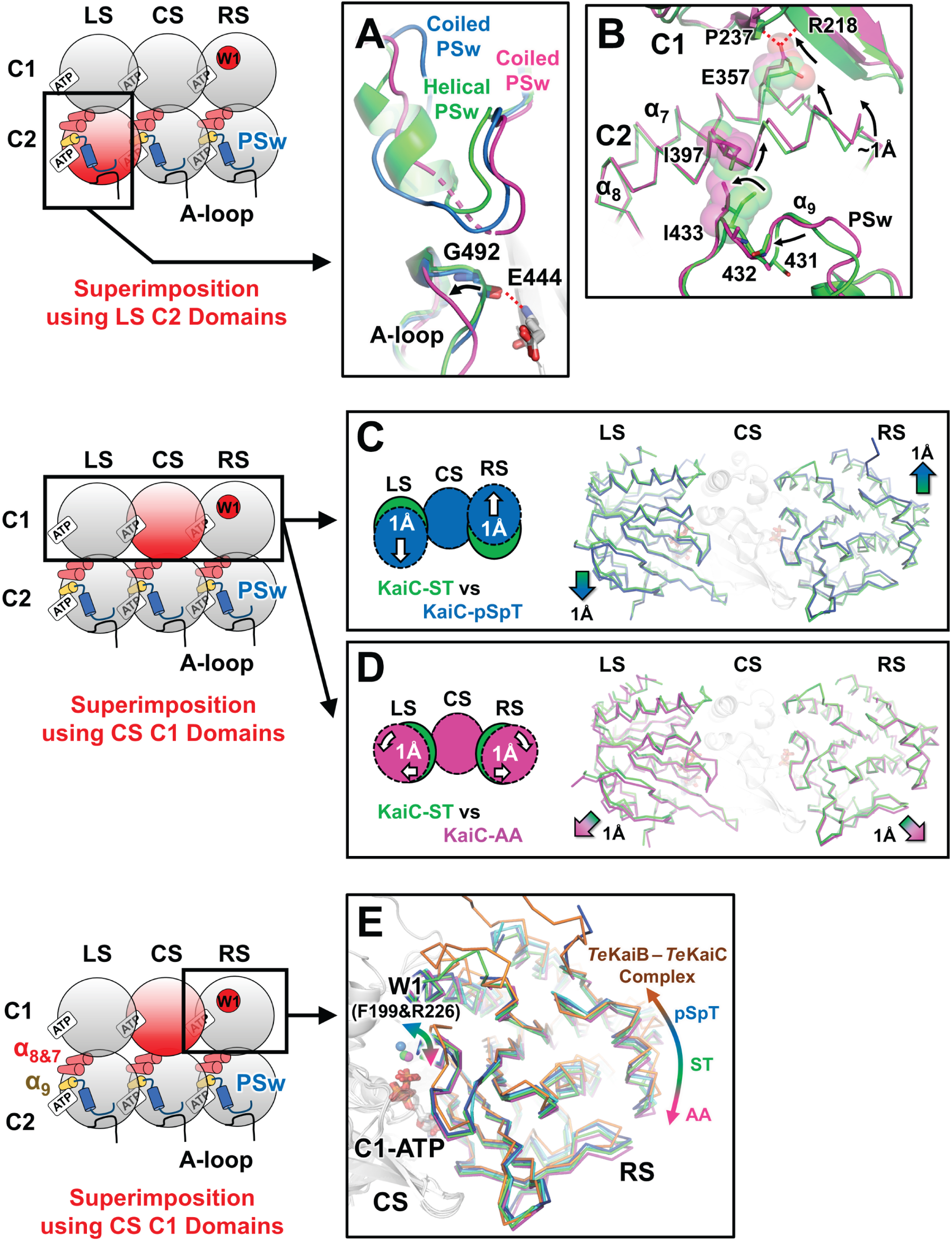
Up- and Downregulation of C1-ATPase via Quaternary Structural Changes. KaiC-ST (green), KaiC-pSpT (blue), and KaiC-AA (magenta) crystal structures are viewed from the inner-radius side of the hexamer and superimposed using a red-colored C1 or C2 domain of *center subunit* (CS), *right subunit* (RS), or *left subunit* (LS) as shown in schematic diagrams. (*A*) Helix-to-coil transition of PSw and unfolding of A-loop in the transition from KaiC-ST to KaiC-AA. The hydrogen bond between G492 and E444 observed in KaiC-ST is disrupted in KaiC-AA. (*B*) C1 ring quaternary structural change assisted by positional shifts of PSw, α_9_, α_8_, and α_7_ in KaiC-AA. α_8_ is lifted up toward the C1 domain by steric contacts between I433 and I397. (*C*) Tilt-and-slide movement of the C1 domain in KaiC-pSpT. (*D*) Outer-radius–directed movement of the C1 domain to form a less compact C1 ring of KaiC-AA. (*E*) Sliding of C1–C1 interface and updating of lytic water (W1) positions. KaiC-pSpT (blue), *Te*KaiC-pSpT (cyan), KaiC-ST (green), KaiC-AA (magenta), and *Te*KaiC-ET (brown) complexed with *Te*KaiB (5JWQ) (30) are superimposed using their C1 domains in the CS subunit. Note that the affinity of KaiC for KaiB is reduced (Fig. 1*E*) as the relative position of the RS-C1 domain to CS-C1 domain shifts in the C2 direction.

PSw of KaiC-AA was found to adopt a loop conformation (Fig. 4*A*), providing the tunnel-shaped void in the C2–C2 interface to allow uptake of bulk water molecules to the C2-ATPase active site (Fig. 3, *C* and *D*). Compared to KaiC-ST, the coiled PSw of KaiC-AA was shifted away from the nearest C2-ATP, similar to what was observed in KaiC-pSpT (Fig. 4*A*). However, the effects of these PSw shifts on two adjacent α-helices (α_8_ and α_7_) were slightly but notably different between KaiC-AA and KaiC-pSpT. In KaiC-AA, the short helix (α_9_) located downstream of PSw was shifted to the outer radius side of the hexamer, and the resultant steric hindrance between I433 (α_9_) and I397 (α_8_) pushed the N-terminus of α_8_ toward the C1 direction by ∼1 Å (Fig. 4*B*). Concomitantly, α_7_ placed in parallel to α_8_ was pushed in the same direction. The side chain of E357 (α_7_) was hydrogen-bonded to P237 and R218 in the C1 domain, and was inserted into the C1 side as α_7_ changed its orientation. These positional shifts of PSw, α_9_, α_8_, and α_7_ in KaiC-AA assist the movement of every C1 domain to the outer-radius side by ∼1 Å (Fig. 4*D*), finally shifting W1 in every C1–C1 interface into a position more suitable for ATP hydrolysis (Fig. 2*D* and 4*E*) and thus further activating C1-ATPase (*t* test: ***significant; p < 0.0005 / 6, **significant; p < 0.005 / 6, Fig. 1*D*).

Coiled and shifted PSw also influenced the C-terminal region of KaiC-AA. Amino acids positioned from 488 to 497 form a loop (A-loop) (24) in both KaiC-ST and KaiC-pSpT, but the loop is deformed in KaiC-AA to allow its KaiA-binding C-terminus (498–519) (25) to be more exposed to solvent (Fig. 4*A*). This observation is consistent with those of previous studies (26, 27) that reported a high affinity of KaiA to KaiC-AA, implying that the structure of KaiC-AA reflects the KaiA-stimulated or KaiA–post-bound state of KaiC-ST. Consistent with the expectation that the A-loop is pulled-down by KaiA binding to the C-terminus of KaiC-ST (24), a key hydrogen bond between G492 and E444 is disrupted in KaiC-AA (Fig. 4*A*).

### Relationship Between the Regulatory State of C1-ATPase^full^ and KaiB Affinity

It has been shown that KaiB binds to the C1 domain of KaiC (28-30), and the affinity and kinetics of KaiB–KaiC interaction have been investigated in detail (31, 32). To evaluate the functional significance of the observed quaternary structural changes of the C1 ring (Fig. 4, *C, D*, and *E*), we studied the affinity of KaiC to KaiB with native polyacrylamide gel electrophoresis (PAGE) analyses. A mixture of equimolar amounts of KaiB and KaiC-ST (5 μM on a monomer basis) after incubation at 30°C for 30 h revealed that 50% of KaiB bound to KaiC-ST (> 97% dephosphorylation) to form a KaiB/KaiC-ST binary complex (*SI Appendix*, Fig S1*A*). Interestingly, as summarized in Fig. 1*E*, the KaiB affinity of a series of KaiC mutants revealed a correlation with the *b*-value; the affinity of KaiC for KaiB decreased as the *b*-value increased (*SI Appendix*, Fig S1*A* and Table S3). This correlation suggests that the C1-ring status responsible for KaiB binding switches in the range of *b*-values between approximately 1.3 and 1.7. While the *b*-value of KaiC-ST was in the transition region, the affinity of KaiC-AA for KaiB was below the present limit of detection by native PAGE, as reported previously (6, 33, 34). Astonishingly, as long as the rules for up- and downregulation are satisfied, KaiC could dramatically switch even from extremely low-affinity (KaiC-AA) to high-affinity forms (KaiC^E357Q^-AA). In this context, KaiC-AA is a poor mimic of KaiC-ST, in terms of not only structure (Fig. 2 and Fig. 3) but also KaiB binding (Fig. 1*E*). Our results clearly indicate that the C1-ATPase^full^ of KaiC-ST is not upregulated enough to completely disrupt the interaction between KaiB and KaiC-ST (Fig. 1*E*).

The complete disassembly of KaiB from KaiC requires the KaiA-assisted activation of the dual ATPase. Immediately after the addition of KaiA (t = 0 h) to the KaiB–KaiC-ST complex that was pre-formed for 48 h (t = −48 to −0.1 h), KaiA was trapped on KaiB interacting with the C1 domain of KaiC-ST (Fig. 5*A*), and then eluted from a size-exclusion chromatography (SEC) column as the KaiA–KaiB–KaiC-ST ternary complex (t = 0, Fig. 5*A*). Within 1.5 h at 30°C, the peak intensity of the ternary complex decreased whereas that of free KaiB increased (inset in Fig. 5*A*), while the dual-phosphorylation site of KaiC-ST remained unchanged (Fig. 5*C*). By contrast, more ADP was produced by the KaiA/KaiB/KaiC-ST mixture than by the KaiB/KaiC-ST mixture within 1.5 h, peaking ∼4 h after KaiA addition (Fig. 5*D*). A small amount of free KaiA (Fig. 5*E* and *SI Appendix*, Fig S1*B*) likely binds the C-terminal loop of KaiC-ST and then activates ATPase to disrupt the interaction between KaiB and KaiC-ST (31), as a KaiC mutant (KaiC^Δ505^) lacking the C-terminus (506–519) (19, 24, 33) was assembled into a KaiA–KaiB–KaiC^Δ505^-ST complex that rarely disassembled (Fig. 5*B*). Notably, the KaiA-assisted disassembly shown in Fig. 5 has the appearance of an autocatalytic reaction often found in oscillators; a KaiA molecule released from a ternary complex can react with another ternary complex to promote the release of another KaiA molecule (*SI Appendix*, Fig. S2*F*).

**Figure 5.**
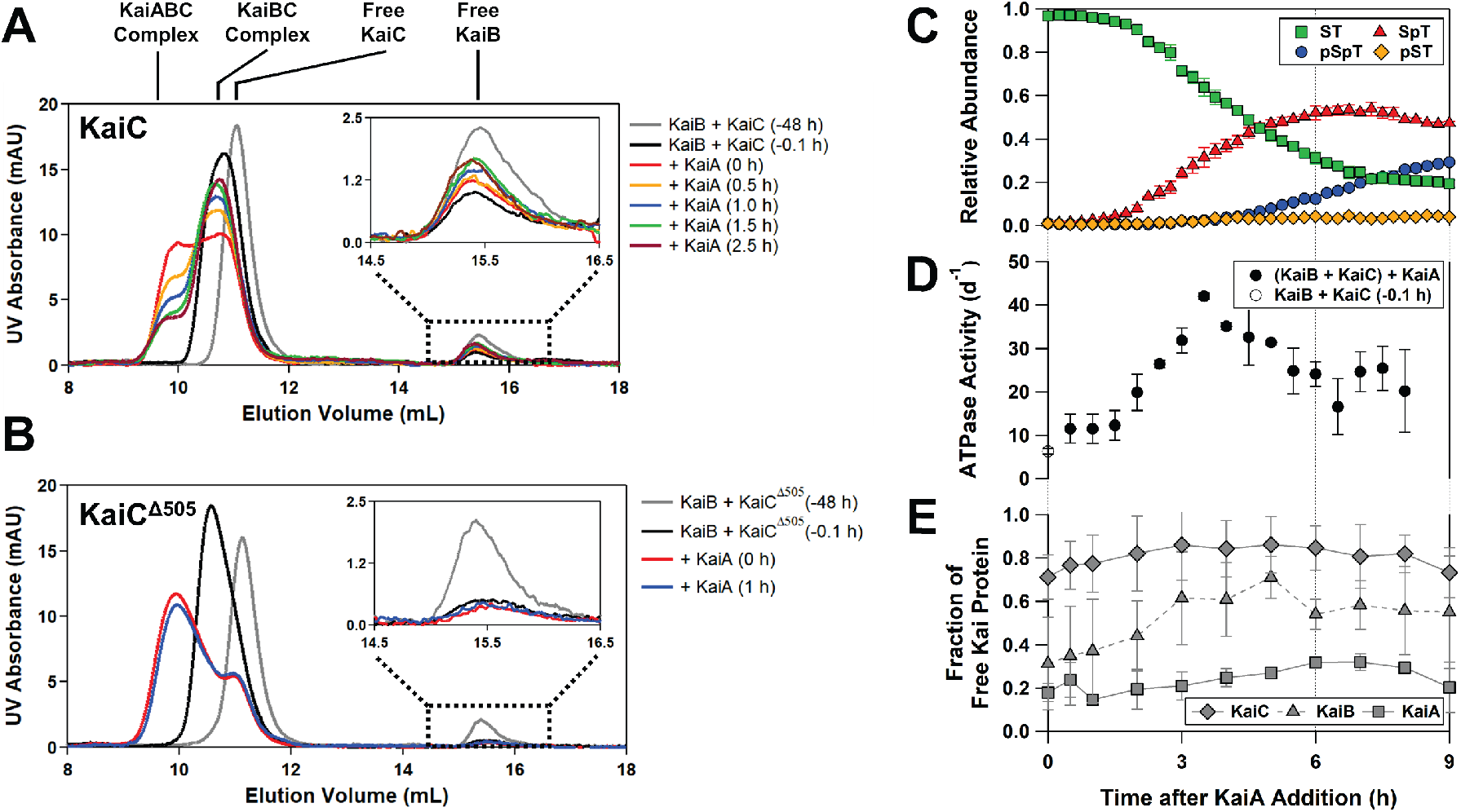
Activated C1/C2-ATPase of KaiC Triggers Disassembly of the KaiABC Ternary Complex. Time evolution of SEC chromatograms before and after adding KaiA to the pre-formed (*A*) KaiB−KaiC-ST complex and (*B*) KaiB−KaiC^Δ505^-ST complex. Black and gray lines correspond to the elution curves of the KaiB/KaiC mixture with and without incubation at 30°C for 48 h, respectively. Other chromatograms show assembly and disassembly dynamics of KaiA−KaiB−KaiC-ST ternary complex after addition of KaiA at t = 0 h. Peak assignments are supported by molecular mass characterization of eluates using multiangle light scattering (*SI Appendix*, Table S4). Inset shows a magnified view of peaks corresponding to free KaiB. Time courses for (*C*) phosphorylation, (*D*) ATPase activity, and (*E*) free Kai proteins after addition of KaiA to the pre-formed KaiB−KaiC-ST complex (*SI Appendix*, Fig S1*B*).

To examine this hypothesis, we conducted a preliminary mathematical model analysis of the KaiA-assisted disassembly. Our present model provides only a qualitative assessment because it does not take into consideration KaiB rebinding to the KaiC hexamer (*SI Appendix*, Extended Method S1). Nevertheless, important aspects of our experimental observations were reflected by the model when optimized parameter values were used. First, the disassembly of the KaiA–KaiB– KaiC-ST complex preceded the accumulation of phosphorylated KaiC (Fig. 5, *A* and *C*, *SI Appendix*, Fig. S2*A*). Second, free KaiB accumulated ∼4 h before free KaiA increased (Fig. 5E, *SI Appendix*, Fig. S2*B*). Third, the concentration of the KaiA–KaiC complex peaked at ∼4 h (*SI Appendix*, Fig. S2*A*) when the time-dependent increase in KaiC-SpT abundance was maximized (Fig. 5*C*). Fourth, the present model (*SI Appendix*, Fig. S2*D*) successfully predicted the KaiA-dependent acceleration of disassembly and phosphorylation dynamics (*SI Appendix*, Fig. S2*E*). Fifth, the KaiA/KaiB/KaiC^Δ505^-ST complex, which was rarely disassembled and phosphorylated (Fig. 5*B*), could be reproduced when the binding rates of KaiA to the C2 domain of KaiC^Δ505^-ST were 30- to 50-fold lower than those to the C2 domain of KaiC-ST (*SI Appendix*, Fig. S2*C*). These findings qualitatively support the KaiA-assisted and auto-catalytic disassembly of the KaiA–KaiB– KaiC-ST complex through the activation of C1/C2-ATPase.

## Discussion

In the presence of KaiA and KaiB, KaiC exhibits circadian rhythms not only in its phosphorylation state (8) but also its C1/C2-ATPase activity (11, 12, 19, 32); C1/C2-ATPase activity was inhibited during the phosphorylating phase and, conversely, was activated during the dephosphorylating phase (Fig. 1*B*). Our biochemical (Fig. 1*C*, *SI Appendix*, Table S3) and structural (Figs. 2 and 3) analyses indicated that both C1-ATPase^full^ and C2-ATPase^full^ were activated or inactivated in a concerted manner (e.g., KaiC-ST vs KaiC-AA), but sometimes under opposing regulation (e.g., KaiC-ST vs KaiC-pSpT).

The present results showed that the properties of KaiC-AA were more similar to those of KaiC-ST stimulated by KaiA than to those of unstimulated KaiC-ST. Even in the absence of KaiA, C1-ATPase^full^ and C2-ATPase^full^ of KaiC-AA were both activated to a similar level as when KaiA was added to KaiC-ST (Fig. 1*C*). In contrast to KaiC-ST, PSw of KaiC-AA was coiled (Fig. 4*A*) to form a tunnel-shaped cavity at the C2–C2 interface for bulk water uptake (Fig. 3*C*), and at the same time, W1 was relocated to a suitable position for ATP hydrolysis (Fig. 2*D*) by the quaternary structural change of the C1-ring (Fig. 4*E*). These results suggest that KaiC-AA retains some of the structural and functional features of KaiC-ST stimulated by KaiA.

Assuming that KaiC-AA can be treated as a functional (Fig. 1*C*) and structural (Fig. 2*D* and 3*D*) mimic of KaiC-ST stimulated by KaiA, we can infer the results of KaiC-ST activation triggered by KaiA by tracing the conformational change from KaiC-ST to KaiC-AA starting from the KaiA binding site (Fig. 6). The folded A-loop hooks the KaiA-binding C-terminus (498–519) of KaiC-ST, which is in a less exposed configuration (Fig. 6*A*), by a hydrogen bond formed between the main-chain carbonyl oxygen atom of G492 and the main-chain amide of E444 (Fig. 4*A*). This sequence of events begins with KaiA binding to the C-terminal tail of KaiC-ST (25) and drawing out the A-loop located on the N-terminal side of the tail (24) (Fig. 6*B*). This induces breaking of the hydrogen bond between G492 and E444, as confirmed in KaiC-AA (Fig. 4*A*). The destabilization of the A-loop causes the helix-to-coil transition of PSw adjacent to the A-loop (Fig. 6*B*). Eventually, the tunnel-shaped void at the C2–C2 interface is expanded and water molecules are taken up through it to the C2-ATPase active site where E318 and C2–ATP are located (Fig. 3). The effect of the structural change in PSw is not confined to the C2 ring but extends to the C1 ring through the C1– C2 interface. The short α_9_ helix downstream of PSw moves along its helical axis to the outer radius side of the hexamer, pushing the α_8_ and α_7_ helices up to the C1 side (Fig. 6*C*). These perturbations cause a change in the quaternary structure of the C1 domain so that the ring structure expands slightly (Fig. 6*D*); the lytic water molecule W1 that reacts with C1–ATP is sequestered at the C1– C1 interface via hydrogen bonds between F199 and R226 (F200 and R227 in *Te*KaiC), and is relocated to a position much more suitable for ATP hydrolysis (C1-ATPase) by the change in the C1–C1 interface.

**Figure 6.**
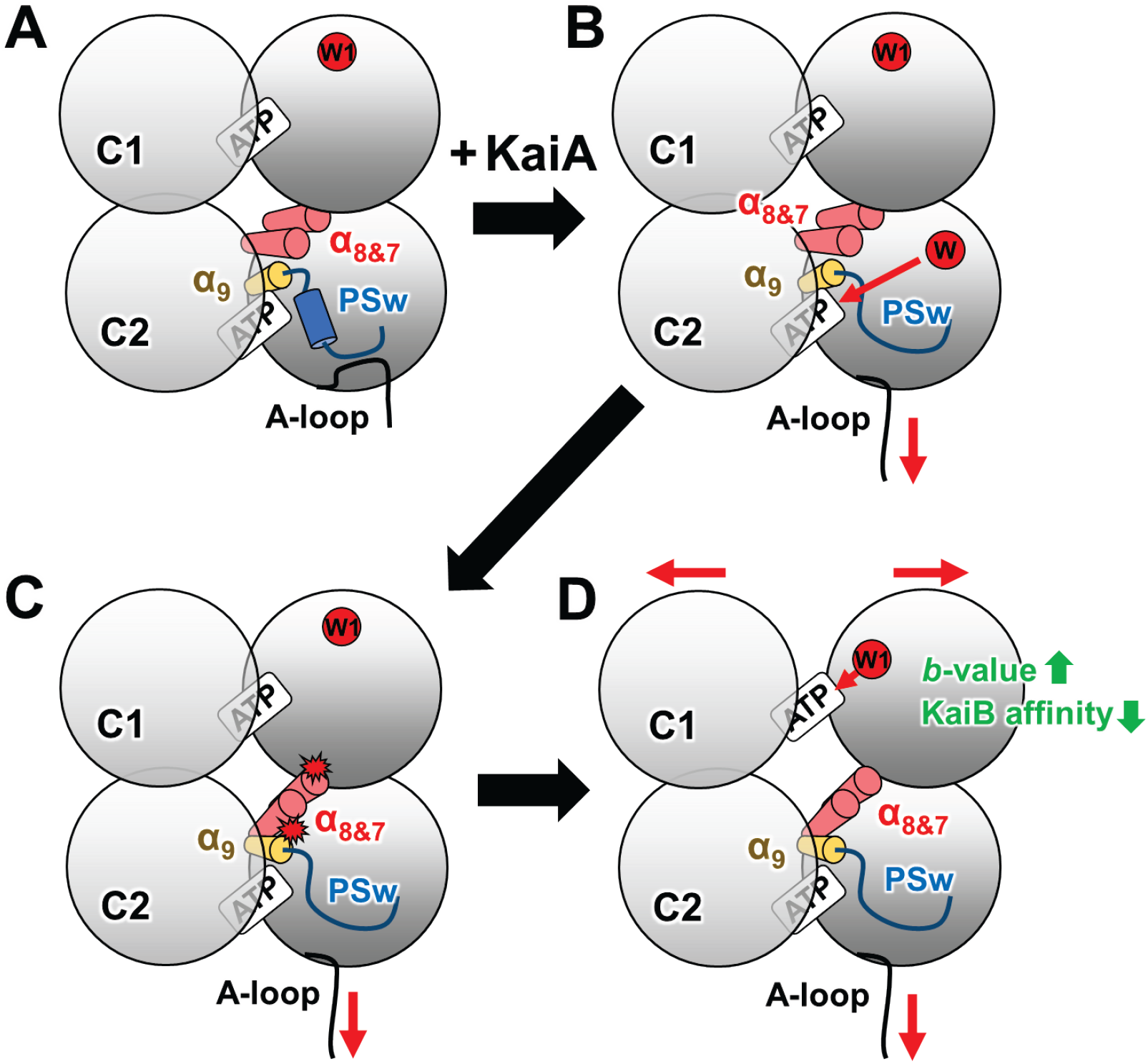
Schematic Drawing of the Activation Mechanism of the Dual ATPase in KaiC-ST. (*A*) KaiC-ST with helical PSw and folded A-loop. A lytic water molecule, W1, is sequestered at a position unfavorable for S_N_2-type C1-ATP hydrolysis. (*B*) Unfolding of the A-loop (24) followed by the helix-to-coil transition of PSw in KaiA-stimulated KaiC-ST. Bulk water molecules, W, migrate into the C2 active site through the tunnel-shaped void in the C2–C2 interface. (*C*) A short α_9_ helix near coiled PSw pushes α_8_ and α_7_ helices up to the C1 side. (*D*) W1 is relocated to a position much more suitable for C1-ATP hydrolysis through the quaternary structural change of the C1 ring.

By contrast, the mechanism of inactivating C1-ATPase will be inferred by tracing the conformational change from KaiC-ST to KaiC-pSpT, or to KaiC-pST (23). The helix-to-coil transition of PSw from KaiC-ST to KaiC-pSpT is coupled to the rearrangement of polar residues constituting electrostatic interactions in the C1–C2 interface (23), which causes the tilt and slide movement of the C1 domains to facilitate the quaternary structural change of the C1 ring (Fig. 4*C*). This update of the C1–C1 interface results in W1 sequestration at a position much less suitable for C1–ATP hydrolysis (Fig. 4*E*).

Our results indicate that the activation of dual KaiC ATPases contributes to a dawn-phase autocatalytic disassembly of the ternary night complexes (Fig. 6, *SI Appendix*, Fig. S2*F*) and is crucial for resetting subjective night and then pushing the system forward along the unidirectional cycle. Although the structural details of C2-ATPase are still unclear because we were unable to identify the lytic water molecules for C2-ATPase, it is simply amazing that it possesses a dynamic range as wide as 0–200 d^-1^ (Fig. 1*C*, *SI Appendix*, Table S3). Given that the activity level of C2-ATPase is suppressed to zero in KaiC-ST, its auto-inhibitory mechanism and its physiological significance are the next important research targets, which will hopefully lead to a better understanding of the structural and functional roles of the KaiC dual ATPase in the circadian clock system.

## Materials and Methods

### Expression, Purification, and Crystallization of KaiC

Plasmid vectors for wild-type KaiC, KaiC mutants, and *Te*KaiC were generated for glutathione S-transferase (GST)-tagged (pGEX-6P-1) (6) or hexahistidine (His)-tagged (pET-3a) (18) forms. Kai proteins were expressed in *E. coli*. BL21(DE3) or BL21(DE3)pLysE and purified as reported previously (18, 35).

All crystals were obtained by the vapor diffusion method as reported previously (23). The *Te*KaiC-pSpT crystal in the *P*2_1_2_1_2_1_ space group was obtained in solutions containing 80 mM acetic acid and 1.0 M sodium acetate, and frozen with 25% (w/v) PEG8K. The KaiC-AA crystal in the *R*3 space group was obtained in solutions containing 100 mM Tris/HCl (pH 7.0), 1 M KCl, 0.8 M sodium/potassium tartrate, 0.95 M sodium acetate, and 5 mM AMP-PNP.

### Data Collection and Structure Determination

Preliminary X-ray diffraction experiments were conducted using FRX-Synergy (RIGAKU, Japan), and the final dataset was collected on beamline BL44XU at SPring-8 (Harima, Japan) at 100 K under a cryostream. Diffraction images were recorded with an MX-300HE (Rayonix) or PILATUS (Eiger) detector and processed using the HKL2000 (36) and XDSGUI (37) software. Initial phases were determined by molecular replacement methods using structures 2GBL (38) or 4O0M (39), and MOLREP (40). Refmac5 (41) and COOT (42) were used to refine and model the crystal structures, respectively. Graphic representations were drawn using PyMOL (Schrödinger). Tunnel-shaped voids were analyzed with a probe radius of 0.6 Å using CAVER 3.0 (43). Statistics of data collection and refinements are listed in *SI Appendix*, Table S1.

### Biochemical Assays of Kai Proteins

The four phosphorylation states of KaiC were detected by SDS-PAGE and then quantified by LOUPE software (44). The number of ATP molecules hydrolyzed into ADP molecules per KaiC monomer per unit time was quantified at 30°C as previously described (18, 19). The ATPase activities were analyzed using one-way ANOVA and Bonferroni post hoc *t* test.

### Determination of *b*-values as a Measure of Up- and Downregulation of C1-ATPase^full^

In this study, the ATPase activity of a truncated KaiC consisting solely of the C1 domain (11) was defined as the basal activity (C1-ATPase^basal^) of C1-ATPase. The ATPase activities of full-length KaiC and its mutants, each of which carries the E318Q substitution that inactivates both kinase and C2-ATPase, were measured to estimate the contribution of C1-ATPase (C1-ATPase^full^) (*SI Appendix*, Table S3). The *b*-value was then calculated as the ratio of C1-ATPase^full^ to C1-ATPase^basal^. The *b*-values were analyzed using one-way ANOVA and Bonferroni post hoc *t* test.

### KaiB–KaiC Interaction Assay

KaiB (0.06 mg/mL) was incubated with KaiC-WT or its mutants (0.3 mg/mL) at 30°C in a buffer containing 20 mM Tris/HCl (pH 8.0), 150 mM NaCl, 0.5 mM EDTA, 1 mM ATP, 5 mM MgCl_2_, and 1 mM DTT. Aliquots were taken from the incubated samples after 30 h and subjected to native PAGE analysis (*SI Appendix*, Fig S1A). Electrophoresis was performed at 25°C using running buffer containing 25 mM Tris/HCl (pH 8.3), 192 mM glycine, 5 mM MgCl_2_, and 1 mM ATP.

### KaiA–KaiB–KaiC Interaction Assay

KaiB and KaiC were mixed in a buffer containing 20 mM Tris/HCl (pH 8.0), 150 mM NaCl, 0.5 mM EDTA, 1 mM ATP, 5 mM MgCl_2_, and 1 mM DTT, and then incubated at 30°C to equilibrate the formation of the KaiB–KaiC-ST complex. After incubation for 48 h, KaiA was added to the equilibrated KaiB/KaiC-ST mixture at a final concentration of 0.06, 0.06, and 0.3 mg/mL for KaiA, KaiB, and KaiC, respectively (1:3:3 molar ratio on a per monomer basis). Every aliquot taken from the KaiA/KaiB/KaiC-ST mixture incubated at 30°C was subjected to SEC and native-PAGE analyses (*SI Appendix*, Fig S1B). A Superdex 200 column (Cytiva) was connected to a multiangle light scattering (MALS) system (Viscotek TDA305, Malvern) to estimate the molecular masses of the eluted peaks (*SI Appendix*, Table S4). Native PAGE analysis was conducted at 25°C using running buffer containing 25 mM Tris/HCl (pH 8.3), 192 mM glycine, 5 mM MgCl_2_, and 1 mM ATP.

## Data Availability

Atomic coordinates and structure factors have been deposited in the Protein Data Bank with the accession codes: 7DY1 (*Te*KaiC) and 7DYE (KaiC-AA).

## Acknowledgments

Diffraction data were collected at BL44XU at the SPring-8 facility under proposals 2017A6700, 2017B6700, 2018A6700, 2018B6700, 2019A6700, 2019B6700, 2020A6700, 2020A6500, 2017A6702, 2017B6702, 2018A6802, 2018B6802, 2019A6902, 2019B6902, and 2020A6502. This research was partly supported by the Platform Project for Supporting Drug Discovery and Life Science Research (BINDS) from AMED under Grant Number JP20am0101072. This study was supported by Grants-in-Aid for Scientific Research (17H06165 to S.A.; 19K16061 to Y.F.; and 18K06171 to A.M.).

