## Supplementary material for "Regulation Mechanisms of the Dual ATPase in KaiC": SI Appendix

### Supplementary Information for
#### Regulation Mechanisms of the Dual ATPase in KaiC

Yoshihiko Furuike[a,b], Atsushi Mukaiyama[a,b], Shinichi Koda[c], Damien Simon[a,b], Dongyan Ouyang[a], Kumiko Ito-Miwa[d], Shinji Saito[c], Eiki Yamashita[e], Taeko Nishiwaki-Ohkawa[f,g], Kazuki Terauchi[h,i], Takao Kondo[d], Shuji Akiyama[a,b,*]

[a]Research Center of Integrative Molecular Systems (CIMoS), Institute for Molecular Science, National Institute of Natural Sciences, 38 Nishigo-Naka, Myodaiji, Okazaki 444-8585, Japan.

[b]Department of Functional Molecular Science, SOKENDAI (The Graduate University for Advanced Studies), 38 Nishigo-Naka, Myodaiji, Okazaki 444-8585, Japan.

[c]Department of Theoretical and Computational Molecular Science, Institute for Molecular Science, National Institute of Natural Sciences, 38 Nishigo-Naka, Myodaiji, Okazaki 444-8585, Japan.

[d]Division of Biological Science, Graduate School of Science and Institute for Advanced Studies, Nagoya University; 464-8602 Nagoya, Japan.

[e]Institute for Protein Research, Osaka University, 3-2 Yamada-oka, Suita 565-0871, Japan.

[f]Graduate School of Bioagricultural Sciences, Nagoya University, 464-8601 Nagoya, Japan.

[g]Institute of Transformative Bio-Molecules (ITbM), Nagoya University, 464-8601 Nagoya, Japan.

[h]Graduate School of Life Sciences, Ritsumeikan University, 1-1-1 Nojihigashi, Kusatsu, Shiga 525-8577, Japan.

[i]College of Life Sciences, Ritsumeikan University, 1-1-1 Nojihigashi, Kusatsu, Shiga 525-8577, Japan.

*Shuji Akiyama


**This PDF file includes:**

Figures S1 to S2
Tables S1 to S4
SI References

#### S.1 Extended Methods: Mathematical Modeling

Our preliminary model for disassembly consists of six species: KaiA dimer ($A_2$), KaiB monomer (B), dephosphorylated KaiC hexamer ($C_6$), phosphorylated KaiC hexamer ($C_6^*$), KaiB–KaiC-ST or KaiA–KaiB–KaiC-ST complex ($A_{2n}B_6C_6$, $0 \le n \le 6$) in which $A_2$ is trapped on B interacting with the C1 domain of $C_6$, and the KaiA–KaiC-ST complex ($C_6A_2$) in which $A_2$ binds the C-terminal tail of $C_6$.

The following three assumptions were made in order to build a simplified model. First, dimeric and tetrameric forms of KaiB were omitted because the monomeric form of KaiB is predominant under the present experimental conditions (1). Second, we assumed cooperative binding of six KaiB monomers to one KaiC hexamer, because KaiB strongly prefers a ring-shaped alignment (1). Third, we assumed immediate disassembly of every complex upon binding of KaiA to its C2 domain, without accumulation of any complexes carrying KaiA on both the C1 and C2 sides.

The model consists of the following reactions:

$$\mathrm{A}_{2n}\mathrm{B}_6\mathrm{C}_6 + \mathrm{A}_2 \xrightarrow{(6-n)k_{1\mathrm{on}}} \mathrm{A}_{2(n+1)}\mathrm{B}_6\mathrm{C}_6 \;\; (0 \le n \le 5), \tag{1}$$

$$\mathrm{A}_{2n}\mathrm{B}_6\mathrm{C}_6 \xrightarrow{n\,k_{1\mathrm{off}}} \mathrm{A}_{2(n-1)}\mathrm{B}_6\mathrm{C}_6 + \mathrm{A}_2 \;\; (1 \le n \le 6), \tag{2}$$

$$\mathrm{A}_{2n}\mathrm{B}_6\mathrm{C}_6 + \mathrm{A}_2 \xrightarrow{k_{2\mathrm{on}1}} (n+1)\mathrm{A}_2 + \; 6\,\mathrm{B} + \mathrm{C}_6 \;\; (0 \le n \le 6), \tag{3}$$

$$\mathrm{C}_6 + \mathrm{A}_2 \xrightarrow{k_{2\mathrm{on}2}} \mathrm{C}_6\mathrm{A}_2, \tag{4}$$

$$\mathrm{C}_6\mathrm{A}_2 \xrightarrow{k_{\mathrm{phos}}} \mathrm{A}_2 + \mathrm{C}_6^*. \tag{5}$$

$k_{1\mathrm{on}}$ of Eq. (1) is the rate constant for the binding of a KaiA dimer to a KaiB–KaiC-ST or KaiA–KaiB–KaiC-ST complex. $k_{1\mathrm{off}}$ of Eq. (2) is the rate constant for the dissociation of a KaiA dimer from a KaiA–KaiB–KaiC-ST complex. In the present model, KaiA binding to the C2 domain proceeds in a stepwise manner as described in Eqs. (3) and (4). First, a KaiA dimer acting on the C2 domain completely disassembles KaiB–KaiC-ST and KaiA–KaiB–KaiC-ST complexes. We denote the rate constant of this process by $k_{2\mathrm{on}1}$. Then, a KaiA dimer binds to the C2 domain of a free KaiC hexamer with a rate constant of $k_{2\mathrm{on}2}$. Finally, $k_{\mathrm{phos}}$ of Eq. (5) represents the rate constant for the phosphorylation of the KaiC hexamer. To focus on the disassembly process of the KaiA–KaiB–KaiC-ST complex, the binding of KaiB to KaiC is included implicitly under the initial conditions but ignored for simplicity after the initiation of the reaction. This simplification is reasonable only in the short time scale of the present simulation, because KaiB binding to KaiC is a relatively slow process that takes as long as ~5 h (2, 3).

The initial conditions for the present simulation were adjusted to mimic the state after addition of the KaiA dimer to the preformed KaiB–KaiC-ST. The total concentrations of KaiA, KaiB, and KaiC on a per monomer basis were set to be 1.7, 5.2, and 5.2 μM, respectively. All KaiC monomers were assumed to be dephosphorylated. Fifty percent of KaiB binds to KaiC (Fig. 5*A*), and 90% of KaiA binds to KaiB that interacts with KaiC in the form of $A_{2n}B_6C_6$ ($0 \le n \le 6$) (Fig. 5*E*), where KaiA is distributed binomially. The remaining quantities of KaiA, KaiB, and KaiC exist in the forms of free $A_2$, B, and $C_6$, respectively.

Values for the rate constants were optimized as described below. First, to reproduce the delayed accumulation of phosphorylated KaiC in the experiment, we defined a loss function by

$$L(\boldsymbol{k}) = \mathrm{C}_6^*(t = 4\ \mathrm{h})^2 + \left[\mathrm{C}_6^*(t = 10\ \mathrm{h}) - \frac{5.2}{6}\right]^2, \tag{6}$$

where $\boldsymbol{k}$ is the set of the rate constants and is obtained by minimizing $L(\boldsymbol{k})$ by the Nelder-Mead method (4). The rate equations of Eqs. (1-5) were integrated by the Runge-Kutta 4$^{\mathrm{th}}$ order method. In this procedure, we fixed $k_{1\mathrm{on}}$ to zero in order to emphasize the delayed accumulation of $C_6^*$. Finally, the resultant parameter values were manually fine-tuned so that the concentrations of $C_6^*$ and other species reflected those in the experimental results. The final parameter values were $k_{1\mathrm{on}}$ = 1.0 μM$^{-1}$ h$^{-1}$, $k_{1\mathrm{off}}$ = 0.1 h$^{-1}$, $k_{2\mathrm{on}1}$ = 3.0 μM$^{-1}$ h$^{-1}$, $k_{2\mathrm{on}2}$ = 5.0 μM$^{-1}$ h$^{-1}$, and $k_{\mathrm{phos}}$ = 0.5 h$^{-1}$.

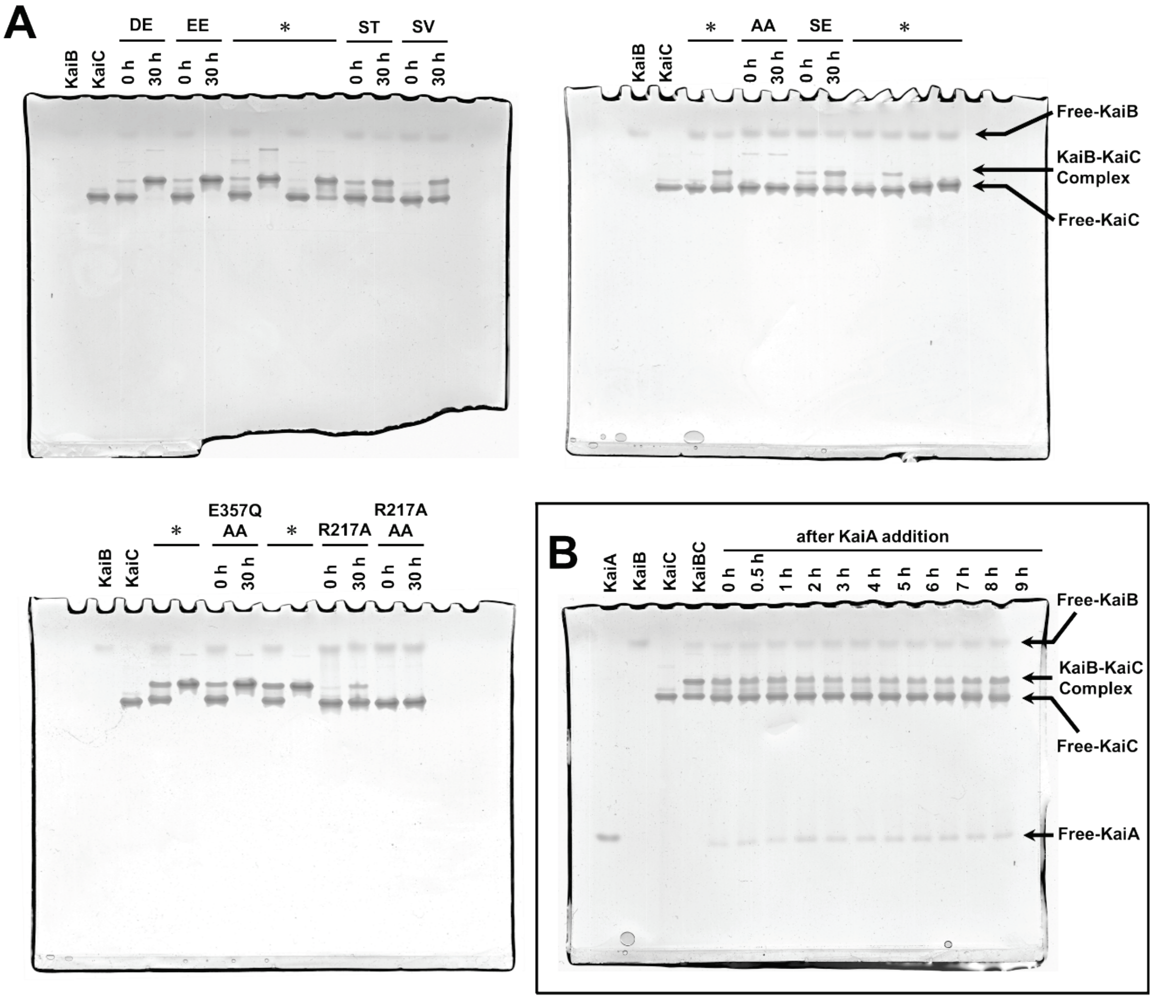


**Fig. S1.** Native PAGE analysis of Kai protein interactions. (*A*) KaiB–KaiC interaction assay at 30°C. Aliquots were taken from the KaiB/KaiC mixture at the indicated times, and then subjected to native PAGE analysis. Bands marked with asterisks are the result of other KaiC mutants not mentioned in this paper. (*B*) KaiA–KaiB–KaiC interaction assay at 30°C. A KaiB/KaiC mixture prepared at t = −48 h was incubated for 48 h to equilibrate the formation of the KaiB−KaiC−ST complex. KaiA was added to the equilibrated KaiB/KaiC-ST mixture at t = 0 h. Every aliquot taken from the KaiA/KaiB/KaiC-ST mixture was subjected to SEC (Fig. 5*A*) and native PAGE analyses. Affinities of Kai protein interactions (Fig. 1*E* and Fig. 5*E*) were estimated by densitometric analysis of gel bands corresponding to free and complexed proteins.

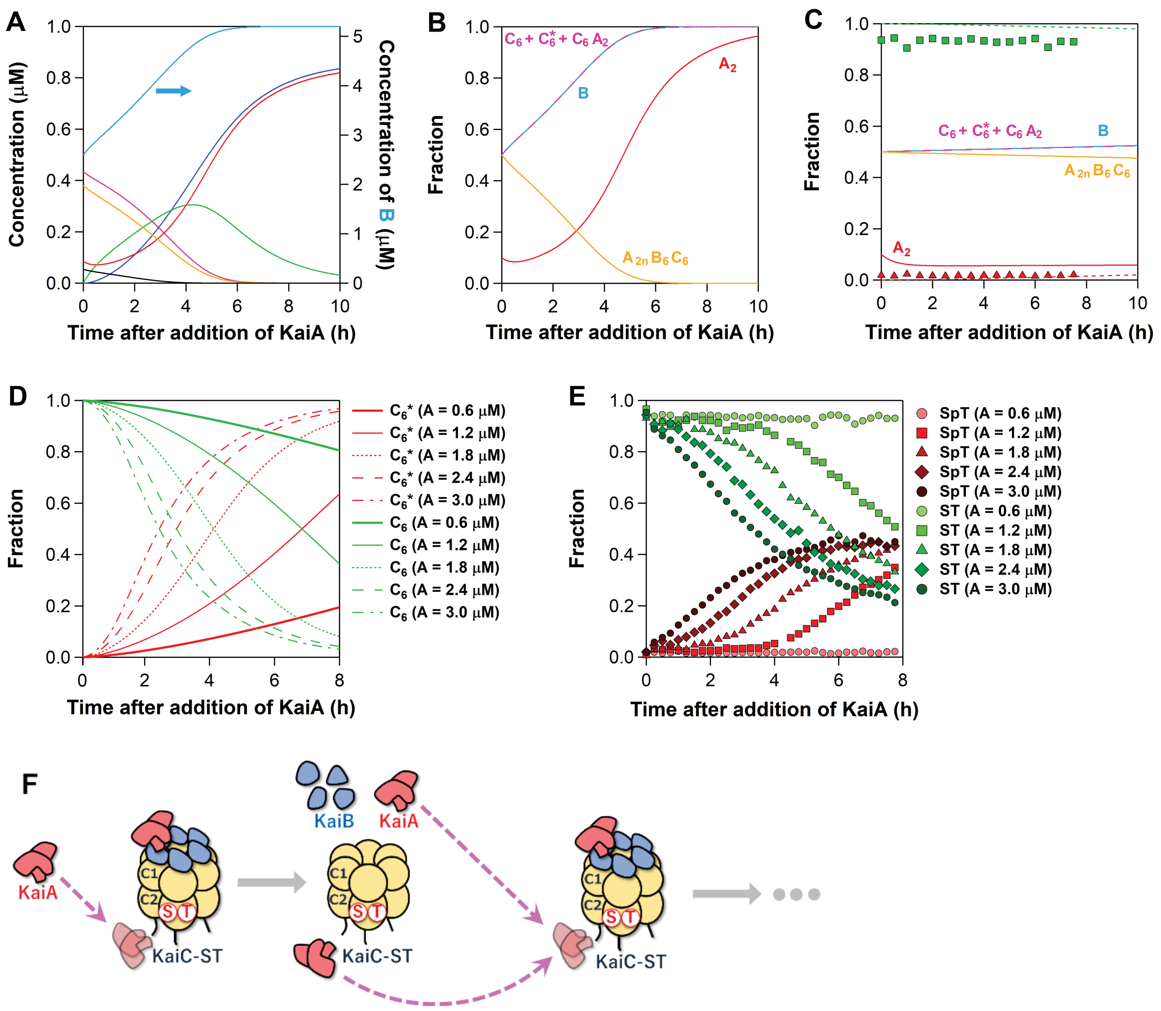


**Fig. S2.** Simulation of KaiA-assisted disassembly model. (*A*) Concentration time courses of KaiA dimer ($A_2$, red line), KaiB monomer (B, cyan line, right axis), dephosphorylated KaiC hexamer ($C_6$, magenta line), phosphorylated KaiC hexamer ($C_6$*, blue line), KaiB–KaiC-ST complex ($B_6C_6$, black line), KaiA–KaiB–KaiC-ST complex ($A_{2n}B_6C_6$: $1 \le n \le 6$, orange line), and KaiA–KaiC-ST complex ($C_6A_2$, green line) after addition of KaiA to the pre-formed KaiB–KaiC-ST complex. (*B*) Time-dependent changes in fractions of night complexes ($A_{2n}B_6C_6$: $0 \le n \le 6$, orange line), $A_2$ (red line), B (cyan dashed line), and the sum of free KaiC hexamer and KaiA–KaiC-ST complex ($C_6$ + $C_6$* + $C_6A_2$, magenta dashed line); in the present native PAGE analysis, the $C_6A_2$ band cannot be separated from that of $C_6$ + $C_6$* (*SI Appendix*, Fig. S1). Note that the amplitude changes in the simulation become larger than those in the experiment (Fig. 5*E*), because KaiB rebinding to the KaiC hexamer is not taken into consideration in the present model. (*C*) The KaiA–KaiB–KaiC$^{\Delta505}$-ST complex, which was rarely disassembled and phosphorylated, could be reproduced under the condition of poorer affinity between KaiA and the C2 domain of KaiC$^{\Delta505}$-ST by using $k_{2on1}$ = 0.1 $\mu M^{-1}$ $h^{-1}$ and $k_{2on2}$ = 0.1 $\mu M^{-1}$ $h^{-1}$. Green and red dotted lines represent simulated fractions of $C_6$ and $C_6$*, respectively. Green squares and red triangles correspond to experimental fractions of KaiC$^{\Delta505}$-ST and KaiC$^{\Delta505}$-SpT, respectively. (*D*) Simulated and (*E*) experimental results of total KaiA-concentration dependence of the KaiC phosphorylation dynamics. Green and red lines (symbols) correspond to the fractions of $C_6$ (KaiC-ST) and $C_6$* (KaiC-SpT), respectively. The total KaiA concentrations are given on a per monomer basis. (*F*) Schematic drawing of a dawn-phase autocatalytic disassembly of the ternary night complexes.

**Table S1.** Data collection and refinement statistics.

| **Protein** | *Te*KaiC-pSpT | KaiC-AA |
|---|---|---|
| **Data Collection** | | |
| Space group | $P2_12_12_1$ | $R3$ |
| Unit cell parameters | | |
| *a, b, c* (Å) | 131.1, 136.5, 190.8 | 94.9, 94.9, 276.5 |
| α, β, γ (°) | 90, 90, 90 | 90, 90, 120 |
| Wavelength (Å) | 0.9 | 0.9 |
| Resolution range (Å)[a] | 47.3–2.20 (2.24–2.20) | 47.5–2.60 (2.72–2.60) |
| Total reflections | 1,219,680 | 150,952 |
| Unique reflections | 172,130 (8248) | 28,555 (3485) |
| Redundancy | 7.1 (7.2) | 5.3 (5.1) |
| Completeness (%) | 99.7 (96.5) | 99.9 (100) |
| $R_{merge}$ (%)[b] | 20.9 (>100) | 6.9 (61.2) |
| (I)/sigma(I) | 6.9 (2.0) | 12.5 (2.8) |
| **Model building** | | |
| Molecular replacement | 4O0M | 2GBL |
| Total atoms | 21,975 | 6,554 |
| Protein | 20,946 | 6,404 |
| Ligands | 192 | 128 |
| Water | 837 | 22 |
| $R_{work}$ (%)[c] | 23.3 | 27.9 |
| $R_{free}$ (%)[c] | 28.4 | 33.7 |
| R.M.S.D. from ideality | | |
| Bond length (Å) | 0.008 | 0.003 |
| Bond angles (°) | 1.5 | 1.2 |
| Average B factors ($Å^2$) | 30.0 | 55.5 |
| Ramachandran plot | | |
| Most favored (%) | 91.3 | 84.9 |
| Allowed (%) | 8.7 | 14.8 |
| Disallowed (%) | 0.0 | 0.2 |
| **PDB code** | 7DY1 | 7DYE |

[a]Values in parentheses are for the highest-resolution shell.
[b]$R_{merge} = \Sigma|I-\langle I\rangle|/\Sigma I$, where I corresponds to the observed intensity of reflections.
[c]$R_{work, free} = \Sigma|F_{obs}-F_{calc}|/\Sigma|F_{obs}|$. $R_{free}$ is the cross-validation of *R*-factor using a subset of randomly selected reflections (5% of the total reflections) that are not included in the refinements.

**Table S2.** Geometric characteristics of lytic water molecules (W1).

| Protein | Subunit | Angle (°) between $O_{3\beta}$, $P_{\gamma}$, and W1 | Distance (Å) between | | |
|---|---|---|---|---|---|
| | | | $P_{\gamma}$ and W1 | F199 O and W1 | R226 $N_{\eta}$ and W1 |
| *Te*KaiC-pSpT‡ | C, D | 137.8, 137.2 | 5.3, 5.0 | 3.9, 3.6 | 3.4, 3.5 |
| | E, F | 143.8, 146.1 | 5.0, 4.9 | 3.4, 3.4 | 3.3, 3.8 |
| | (average) | (141.2) | (5.1) | (3.6) | (3.5) |
| KaiC-ST# | A, B | 149.2, 143.7 | 3.6, 4.3 | 2.7, 3.0 | 2.7, 3.6 |
| | (average) | (146.5) | (4.0) | (2.9) | (3.2) |
| KaiC-AA‡ | A | 157.3 | 3.4 | 2.9 | 2.8 |

‡, *Te*KaiC (7DY1) and KaiC-AA (7DYE) are determined in this work. #, previously deposited structure (7DYJ) (5).

**Table S3** Biochemical analysis of C1/C2-ATPase activities and *b*-values for KaiC and its mutants.

| | C1/C2-ATPase Activity ($d^{-1}$) | | | | | | | | | | | | | | | | | | | *b* value | | | | | Fraction of KaiB in KaiB-KaiC Complex |
|---|---|---|---|---|---|---|---|---|---|---|---|---|---|---|---|---|---|---|---|---|---|---|---|---|
| | (-) E318Q | | | | | (+) E318Q | | | | | C1-ATPase$^{basal}$ | | | | | C1-ATPase$^{full}$ | | C2-ATPase$^{full}$ | | | | | | | |
| | Individual values | | | Ave. | S.D. | Individual values | | | Ave. | S.D. | Individual values | | | Ave. | S.D. | Ave. | S.D. | Ave. | S.D. | Individual values | | | Ave | S.D. | |
| KaiC-ST | 13.9 | 14.9 | 12.1 | 13.6 | 1.4 | 14.6<br>15.2<br>15.7<br>16.1 | 16.6<br>15.0<br>16.2<br>16.5 | 15.6<br>16.5<br>16.2<br>16.0 | 15.8 | 0.6 | 10.0* | 10.2* | 10.9* | 10.4* | 0.5* | 15.8 | 0.6 | -2.2 | 1.6 | 1.40<br>1.46<br>1.51<br>1.55 | 1.60<br>1.44<br>1.56<br>1.59 | 1.50<br>1.59<br>1.56<br>1.54 | 1.52 | 0.10 | 0.45 |
| KaiC-ST + KaiA | 33.6 | 33.3 | 33.3 | 33.4 | 0.2 | 16.8<br>20.7<br>16.8<br>17.0 | 18.8<br>20.0<br>17.1<br>16.5 | 17.8<br>21.4<br>17.9<br>17.8 | 18.2 | 1.7 | | * | | * | * | 18.2 | 1.7 | 15.2 | 1.7 | 1.62<br>1.99<br>1.62<br>1.63 | 1.81<br>1.92<br>1.64<br>1.59 | 1.71<br>2.06<br>1.72<br>1.71 | 1.75 | 0.18 | n.d. |
| KaiC-EE | 5.1 | 6.3 | 6.8 | 6.1 | 0.9 | 2.2 | 2.0 | 1.8 | 2.0 | 0.2 | | * | | * | * | 2.0 | 0.2 | 4.1 | 0.9 | 0.21 | 0.19 | 0.17 | 0.19 | 0.02 | 1.00 |
| KaiC-AA | 22.0<br>21.8 | 28.6<br>22.3 | 26.4<br>22.7 | 24.0 | 2.8 | 19.2 | 18.7 | 19.1 | 19.0 | 0.3 | | * | | * | * | 19.0 | 0.3 | 5.0 | 2.9 | 1.85 | 1.80 | 1.84 | 1.83 | 0.09 | 0.00 |
| KaiC$^{E357Q}$-AA | 5.3 | 5.0 | 5.0 | 5.1 | 0.2 | 11.5 | 11.0 | 12.3 | 11.6 | 0.7 | | * | | * | * | 11.6 | 0.7 | -6.5 | 0.7 | 1.11 | 1.06 | 1.18 | 1.12 | 0.08 | 1.00 |
| KaiC$^{R217A}$ | 21.1 | 17.4 | 20.2 | 19.6 | 1.9 | 11.3 | 11.1 | 10.8 | 11.1 | 0.3 | 8.5 | 8.9 | 9.6 | 9.0 | 0.6 | 11.1 | 0.3 | 8.5 | 1.9 | 1.26 | 1.23 | 1.20 | 1.23 | 0.08 | 0.30 |
| KaiC$^{R217A}$-AA | 237.8 | 217.9 | 186.9 | 214.2 | 25.7 | 19.6 | 19.0 | 19.1 | 19.2 | 0.3 | 8.5 | 8.9 | 9.6 | 9.0 | 0.6 | 19.2 | 0.3 | 195.0 | 25.7 | 2.18 | 2.11 | 2.12 | 2.14 | 0.14 | 0.00 |
| KaiC-SV | 14.7 | 14.8 | 16.0 | 15.2 | 0.7 | 15.1 | 14.5 | 16.6 | 15.4 | 1.1 | | * | | * | * | 15.4 | 1.1 | -0.2 | 1.3 | 1.45 | 1.39 | 1.60 | 1.48 | 0.13 | 0.45 |
| KaiC-DE | 11.4 | 9.6 | 10.6 | 10.5 | 0.9 | 3.6 | 3.6 | 3.8 | 3.7 | 0.1 | | * | | * | * | 3.7 | 0.1 | 6.9 | 0.9 | 0.35 | 0.35 | 0.37 | 0.35 | 0.02 | 1.00 |
| KaiC-SE | 22.0 | 22.7 | 22.6 | 22.4 | 0.4 | 14.2 | 14.4 | 15.2 | 14.6 | 0.5 | | * | | * | * | 14.6 | 0.5 | 7.8 | 0.7 | 1.37 | 1.38 | 1.46 | 1.40 | 0.08 | 0.34 |

*, C1-ATPase$^{basal}$ of KaiC mutants without mutations in the C1 domain were all set to the ATPase activity of a truncated KaiC (KaiC1) consisting solely of the C1 domain (6).

**Table S4** Molecular mass characterization of the Kai protein complexes using SEC-MALS.

| Sample | Elution volume (mL) | | | |
|---|---|---|---|---|
| | 10.0 | 10.7 | 11.0 | 15.4 |
| KaiB + KaiC (−48 h) | n.d. | n.d. | 329 | 52 |
| KaiB + KaiC (−0.1 h) | n.d. | 370 | n.d. | 50 |
| + KaiA (0.0 h) | 535 | 340 | n.d. | 48 |
| + KaiA (0.5 h) | 538 | 346 | n.d. | 50 |
| + KaiA (1.0 h) | 547 | 347 | n.d. | 46 |
| + KaiA (1.5 h) | 554 | 349 | n.d. | 50 |
| + KaiA (2.5 h) | 553 | 349 | n.d. | 47 |
| Assembly State | KaiA–KaiB–KaiC | KaiB–KaiC | KaiC Hexamer | KaiB Tetramer |

The averaged molecular masses of eluted peaks shown in Fig. 5*A* are indicated in kDa. The theoretical molecular masses of KaiC hexamer, KaiB tetramer, and KaiA dimer are 348, 44.7, and 65.3, respectively. Note that the estimated molecular masses for the peaks at 10.0 mL exceed the maximum value for the KaiB–KaiC binary complex (415 kDa).
